# Abrupt transitions to tumor extinction: A phenotypic quasispecies model

**DOI:** 10.1101/044644

**Authors:** Josep Sardanyés, Regina Martínez, Carles Simó, Ricard Solé

**Affiliations:** CREA-Complex Systems Lab, Department of Experimental and Health Sciences, Universitat Pompeu Fabra, Dr. Aiguader 88, 08003 Barcelona, Spain; Institut de Biologia Evolutiva (CSIC-Universitat Pompeu Fabra), Passeig Maritim de la Barceloneta 37, 08003 Barcelona, Spain; Departament de Matem`atiques i Inform`atica (Universitat de Barcelona), Gran Via de les Corts Catalanes 585, 08007 Barcelona, Spain; Departament de Matem`atiques, Edifici C. Facultat de Ci`encies (Universitat Aut`onoma de Barcelona), 08193 Bellaterra, Spain; Santa Fe Institute, 1399 Hyde Park Road, 87501 Santa Fe (NM), USA

**Keywords:** Applied mathematics, Cancer evolution, Cancer targeted therapy, Dynamical systems, Genomic instability, Phenotypic model, Quasispecies

## Abstract

**Background:** The dynamics of heterogeneous tumor cell populations competing with healthy cells is an important topic in cancer research with deep implications in biomedicine. Multitude of theoretical and computational models have addressed this issue, especially focusing on the nature of the transitions governing tumor clearance as some relevant model parameters are tuned. In this contribution, we analyze a mathematical model of unstable tumor progression using the quasispecies framework. Our aim is to define a minimal model incorporating the dynamics of competition between healthy cells and a heterogeneous population of cancer cell phenotypes involving changes in replication-related genes (i.e., proto-oncogenes and tumor suppressor genes), in genes responsible for genomic stability, and in house-keeping genes. Such mutations or loss of genes result into different phenotypes with increased proliferation rates and/or increased genomic instabilities. Also, lethal phenotypes with mutations or loss of house-keeping genes are present in our model.

**Results:** Despite bifurcations in the classical deterministic quasispecies model are typically given by smooth, continuous shifts (i.e., transcritical bifurcations), we here identify an novel type of abrupt transition causing tumor extinction. Such a bifurcation, named as *trans-heteroclinic*, is characterized by the exchange of stability between two distant fixed points (that do not collide) involving, respectively, tumor persistence and tumor clearance. The increase of mutation and/or the decrease of the replication rate of tumor cells involves this catastrophic shift of tumor cell populations. The transient times near bifurcation thresholds are also characterized, showing a power law dependence of exponent –1 of the transients as mutation is changed near the bifurcation value.

**Conclusions:** An abrupt transition involving tumor clearance has been identified with a phenotypic quasispecies cancer model. This result is discussed in the context of targeted cancer therapy as a possible therapeutic strategy to force a catastrophic shift by delivering mutagenic and cytotoxic drugs inside tumor cells. Our model also reveals a novel mechanism causing a discontinuous transition given by the stability exchange of two distant fixed points, which we name as a *trans-heteroclinic* bifurcation.

## Background

A major issue in dealing with cancer progression is the heterogeneous nature of their populations, resulting from genetic instability. This is a defining feature of most advanced tumors [1, 2]. A major consequence of this lack of clonal structure is an enhanced potential to evade cell proliferation checkpoints. Genomic instability refers to an increased tendency of alterations in the genome during the life cycle of cells. Normal cells have a mutation rate of about 1.4 × 10^−10^ changes per nucleotide and replication cycle. It has been proposed that the spontaneous mutation rate in normal cells is not sufficient to account for the large number of mutations found in human cancers. Indeed, studies of mutation frequencies in microbial populations, in both experimentally-induced stress and clinical cases, reveal that mutations that inactivate mismatch repair genes result in 10^2^ — 10^3^ times the background mutation rate, with comparable increases in cancer cells [3–11]. The presence of high instability creates a peculiar situation, since cancer cells become less differentiated, more plastic to adapt but also more prone to failure. How much instability can be afforded by cancer cells? It has been suggested that unstable cancer progression is feasible up to some critical instability levels [15,16]. Once this critical point is reached, instability levels become lethal.

Evidence suggests that cancer populations might actually evolve towards the critical instability point [18]. Such evolution towards the edge of instability has been reported in RNA viruses, where thresholds of lethal mutagenesis have been found to exist [24,25,40]. RNA viruses exhibit critical mutation rates [19,20], beyond which they can experience extinction and their heterogeneous populations are known as *quasispecies* [21] and share several key features with cancer cell populations [16,18,22]. Following these similarities, it was suggested that cancer might also share the presence of critical instability levels [13,15,16,27]. If true, cancer treatments could incorporate increased mutagenesis that would push the system slightly beyond its critical limits [7]. In this context a possible therapeutic strategy in tumors would benefit from targeting DNA repair pathways [28-31]. Similarly, germline mutations in the proofreading domains of DNA polymerases Pol *δ* and *ϵ* have been identified in many types of cancers, giving place to the so-called ‘ultramutator’ phenotype [32]. It has been suggested that increased mutation in these tumors (e.g., anaplastic astrocytoma, breast cancer, or colorectal cancer, among others) with impaired exonuclease proof-reading activities could drive cancer towards the critical transition given by the error threshold [32].

Theoretical models on cancer quasispecies have revealed smooth, continuous transitions towards tumor impairment or collapse as some of the model parameters (typically mutation rates) are tuned (see, e.g., [15,16,26,27,33]). These models deal with mutation rates as single parameters that can be tuned to explore their impact on tumor progression. However, a richer and more useful framework should include a repertoire of changes affecting not only stability but also replication traits. Similarly, the limits imposed to instability levels are largely determined by the presence of essential (house-keeping) genes whose integrity needs to be preserved [18]. A digital genome model incorporating multiple genes associated to replication and stability, as well as the use of house-keeping genes suggests that a complex evolution unfolds as the cancer population approaches higher, near-critical instability levels [17,18]. However, an analytical approach to this more complete picture is still lacking.

Here we study this problem by means of a phenotypic quasispecies model. Our model is similar to a phenotypic model that has been recently used to investigate the effects of mutational fitness effects in RNA viral populations [34]. In particular, we consider a mean field model describing the evolutionary dynamics of different tumor cell phenotypes competing with healthy cell populations. The tumor phenotypes are the result of mutations in proto-oncogenes, tumor suppressor genes, and house-keeping genes, driven by genetic instability. We will specifically model how genomic instability and tumor cells proliferation affect the competition dynamics. To do so we will analyze a minimal model of phenotypic quasispecies, providing the equilibrium points and their stability dependence on model parameters. We will also investigate how the mutation rate of tumor cells, and both genomic instability and proliferation rates affect the transients towards tumor persistence and tumor extinction. Our results will be discussed in the context of the current development of targeted cancer therapies.

## Model and Methods

### Mathematical model

To analyze the dynamics of unstable cancer progression we build a mathematical model using Eigen’s quasispecies model [19], following previous modeling on phenotypic quasispecies [34]. The quasispecies model can be represented, in its general form, by:

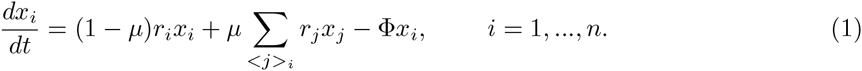

This model describes the time dynamics of a population of *n* sequences with concentration *x_i_* under the processes of replication (proportional to replication rate *r_i_*) and mutation (proportional to mutation rate *µ*) within the assumption of a constant population (CP). The first RHS term denotes the error-free replication of species *x_i_*, while the second RHS term is the synthesis of sequence *i* by mutation from mutant neighbours (denoted as < *j* >*_i_*) in sequence space. Finally, the last RHS term is the dilution flow, which keeps a CP, and is introduced with function Φ, which depends on *x_i_*. The dilution term introduces competition between all *n* sequences (see below for the computation of this term).

Equation (1) will be used in the present manuscript to model the competition between healthy and tumor cells with different replication rates, mutation rates, and survival properties given by the bits composition of sequence *i* (see Fig. 1A,B). Hence, each cell *x_i_* has a genetic state *i* defined as a 3-bits sequence, where *i* denotes the decimal number of the binary sequence. Each bit denotes a different genomic compartment containing replication-related (*R*) genes, genes involved in genomic stability (*S*), and house-keeping (*H*) genes. It is known that alterations in these genes are responsible for tumorigenesis [12]. Hence, such sequences provide the phenotypic traits of the cells of the population. Healthy cells have sequences 000, since no mutations or genomic aberrations are found in these compartments. Tumor cells can have mutations or other types of genomic aberrations such as gene loss in any of these three compartments [12], such abnormalities will be denoted with bit 1.

**Figure 1.**
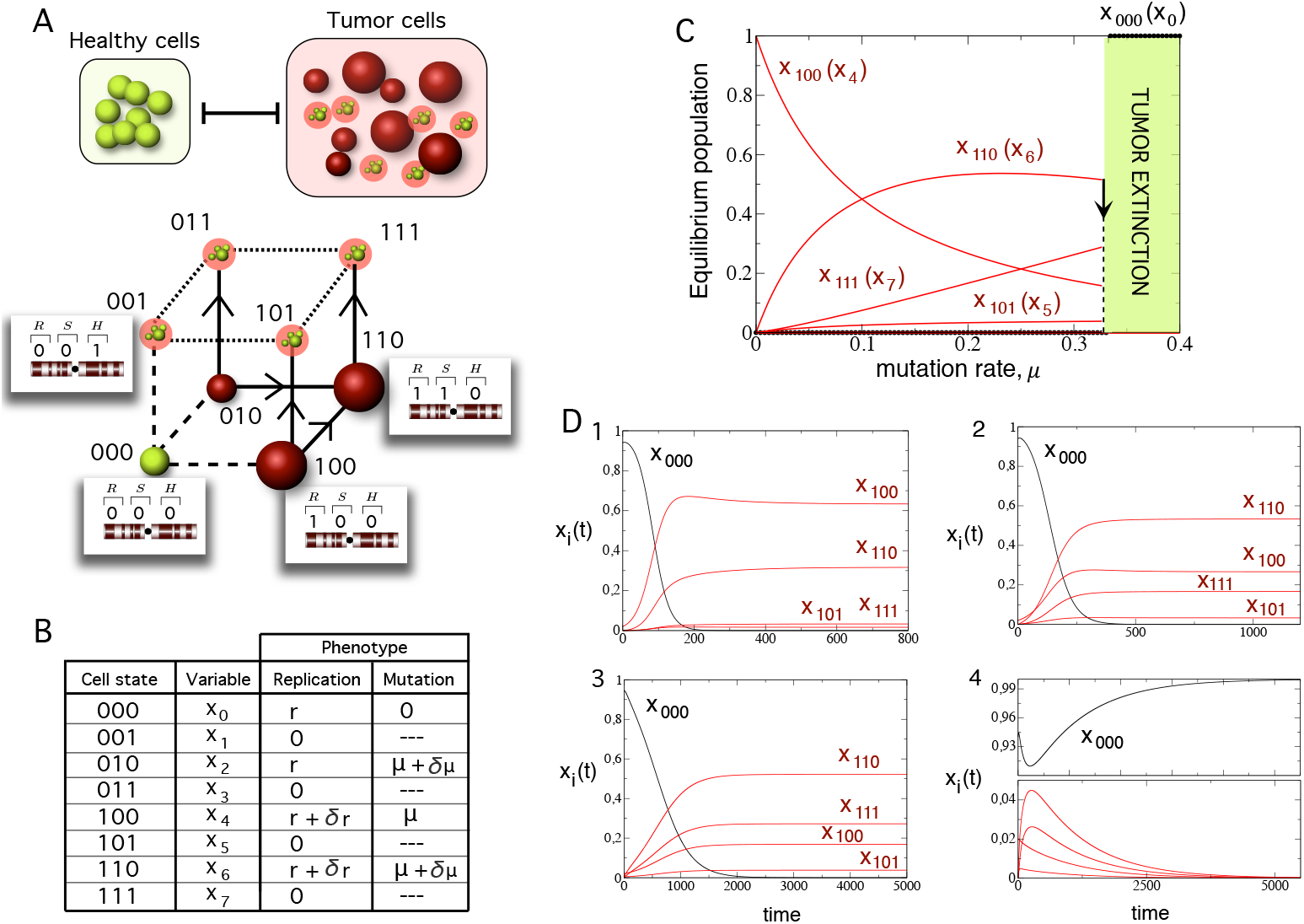
Population structure of the phenotypic quasispecies considering competition between healthy and tumor cells. (A) Our model describes the dynamics of healthy cells (green) competing with a pool of heterogeneous tumor cell phenotypes (red). Each cell is described by a binary state containing three compartments with genes involved in replication (R, such as proto-oncogenes or tumor-suppressor genes), genome stability (S), and the house-keeping (H) genes. The sequence of this cell state determines its phenotypic traits. The system is a disconnected cube since mutations in healthy cells are neglected (dashed lines). Notice that genomes with a mutation or alterations in the H compartment are not viable and do not produce mutant cells (dotted lines). (B) Table displaying (by columns): the binary sequence of each cell; the variable with the subindex corresponding to the integer number of each binary sequence; the replication rate of each cell; and the mutation rate of each genome. (C) Bifurcation diagram using mutation rate of tumor cells, *µ*, as control parameter using *r* = 0.1 and *δ_µ_* = *δ_r_* = 0.05. Notice that the bifurcation is discontinuous as the control parameter *µ* crosses its bifurcation value. (D) Time series using same parameter values as in (C) using: Panel 1: *µ* = 0.05; Panel 2: *µ* = 0.2: Panel 3: *µ* = 0.31; and Panel 4: *µ* = 0.35. The time dynamics of healthy and tumor cells are displayed in black and in red, respectively.

Mutations or genes loss in compartment *R* involve the impairment of genes affecting the rate of replication of cells, which usually increase cells replication activity (introduced in our model with parameter *δ_r_*, see below). This set would include both anomalies in tumor suppressor genes (e.g., APC or P53) and proto-oncogenes (RAS or SRC). Although they act in different ways (are targeted in opposite ways by genetic alterations) here we make no explicit distinction [12]. This assumption is based on the fact that, in terms of population dynamics, alterations in both types of genes lead to a neoplastic process through increases in cancer cells numbers. Mutations in replication-related genes that confer an increase of fitness and thus a selective advantage are named driver mutations. Mutations or genes loss in compartment *S* involve the impairment of genomic stability genes [1,12], which are typically genes playing a key role in preserving genome integrity. These genes (including e.g., BRCA1, BLM, ATM, or P53) keep genetic changes under control. Abnormalities in this compartment will involve an increased level of mutation or genes loss rates, parametrized in our model with *δ_µ_* (see below). Finally, the cells in our model also include a compartment for the so-called house-keeping (*hk*) genes. These genes are tied to essential functions whose failure leads to cell death. In real cells, *hk* genes are constitutive and would include, for example, ribosomal proteins, glyceraldehyde-3-posphate dehydrogenase (GAPDH), or ubiquitin [35], among others.

For our system, as we mentioned above, the state variables *x_i_* are the population numbers of cells with sequence *i*. Parameter *µ_i_* is the mutation rate or genes loss rate of cell *x_i_*, and *r_i_* is the replication rate of the cell with the *i*-th sequence. Finally, Φ is a population constraint which introduces competition between cells also ensuring that the total population remains constant (see below). The full model obtained from Eq. (1), considering the particular structure of the sequence space of our system (see Figure 1A), is given by:

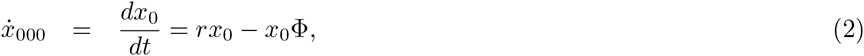

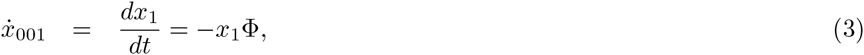

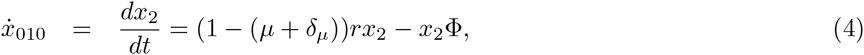

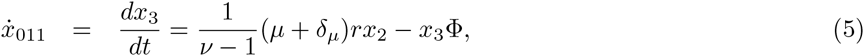

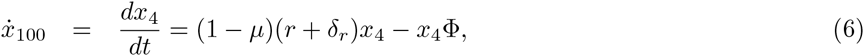

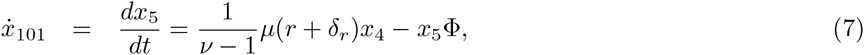

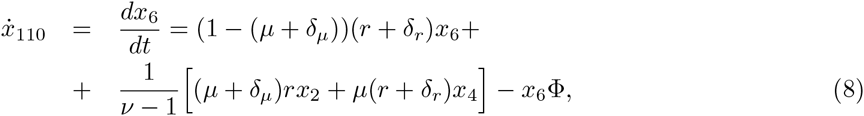

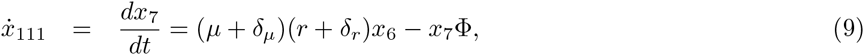

with *ν* = 3. Equation (2) describes the dynamics of healthy cells, which compete with tumor cells. Equation (3) corresponds to the dynamics of a lethal phenotype with cell state 001. Also, Equations (5), (7), and (9) describe the dynamics of lethal phenotypes. All lethal phenotypes have mutations or loss of genes in the house-keeping genes compartment, and are not able to replicate. Hence, they can only be originated from tumor cells able to replicate. Equation (4) corresponds to the dynamics of tumor cells with the mutations or alterations in the stability compartment, giving place to a genomically unstable phenotype. Equation (6) describes the dynamics of tumor cells with mutations or genomic aberrations in replication-related genes compartment. Such phenotype acquires increased proliferation rates. Finally, another possible phenotype can contain both alterations in replication- and stability-related genes compartments. The dynamics of this phenotype is modeled with Eq. (8). Recall that variables subindices are the binary sequence expressed in integers.

Parameter *r* is the basal replication rate of all cells. As mentioned, mutations or aberrations in compartment *R* increase the proliferation rate of tumor cells (parametrized with *δ_r_*). Parameter *µ* is a background mutation rate of tumor cells. Mutation in healthy cells is set to zero since we assume that it is negligible compared to the mutation rate of tumor cells. Anomalies in compartment *S* will increase the mutation of tumor cells by means of parameter *δ_µ_* (e.g., sequences 110 and 010). The modeled system spans a disconnected cube where mutations only affect tumor cells and healthy cells are only coupled to the other variables due to competition (see Table in Fig. 1 for a complete description of the system). As mentioned above, the term Φ is a dilution flow that is used to keep a CP constrain. Hence, by setting 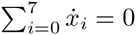, and 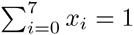, we obtain 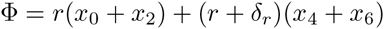.

#### Numerical tools

Numerical solutions have been obtained with the adaptive Runge-Kutta-Fehlberg algorithm of seventh-eighth order, with an automatic step size control fixing the local relative tolerance to 10^−15^.

## 1 Results and Discussion

Firstly, we will characterize the fixed points of the system Eqs. (2)-(9). Then we will determine their stability and the possible bifurcations separating tumor persistence from tumor clearance as a function of the model parameters. Also, we will compute explicit analytical solutions of the model. Finally, we will investigate the transients times near bifurcation thresholds. All of the analytical calculations will be complemented by means of numerical simulations to illustrate the most relevants results in terms of cancer biology and therapy.

### 1.1 Fixed points and stability analysis

The model parameters are given by 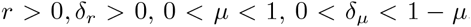. For the sake of simplicity we introduce 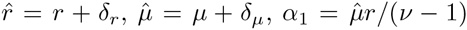, and 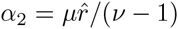. Since *ν* = 3, we have 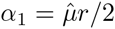, and 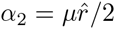. According to the previous terms, and after rearranging the equations, the model under investigation can be represented as follows

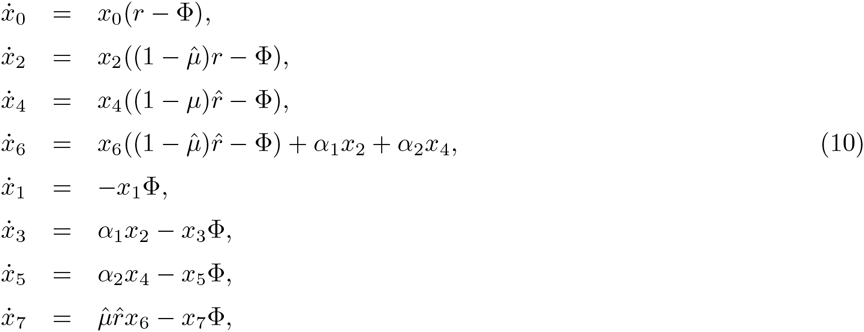

where 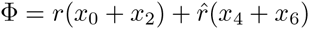. We are interested in values *x_j_* ≥ 0 for all the *x_j_* variables.

It is well known [36, 37] that this kind of systems can be reduced to a linear one with constant coefficients and so, the solutions can be obtained explicitly. However from a dynamical point of view it is also interesting to know the equilibrium points as well as their stability properties.

First, we look for the equilibrium points of (10). If we denote them by (*ξ*, *η*) where *ξ* = (*x*_0_, *x*_2_, *x*_4_, *x*_6_), and *η* = (*x*_1_, *x*_3_, *x*_5_, *x*_7_), it is easy to obtain the following equilib-ria

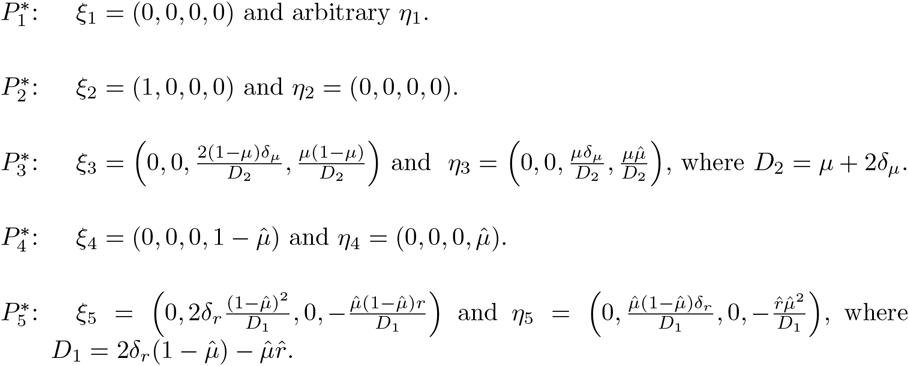

Equilibria 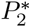 involves, if stable, the dominance of healthy cell populations and the clearance of tumor populations. On the contrary, equilibrium point 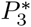 involves, also if stable, the out competition of healthy cells by the tumor cell populations. These two equilibrium points are thus the most relevant ones since they represent an asymptotic extinction and persistence of tumor cells, respectively. Moreover, two lines of equilibria appear for special values of the parameters:

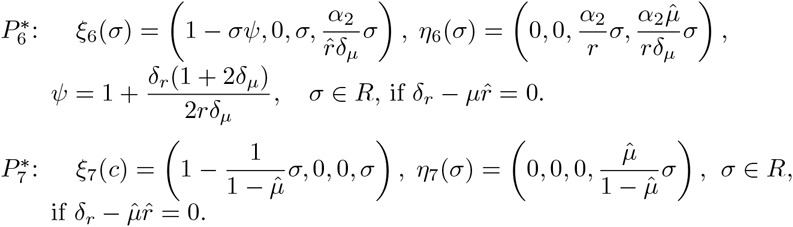

We note that all the components of the equilibria 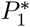 to 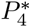 are non negative but for the equilibrium 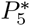 the values of *x*_2_ and *x*_6_ have opposite sign and, hence, it will be discarded. If 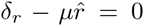 the equilibria 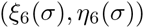 have non negative components for 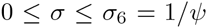. It is easy to check that 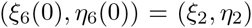 and 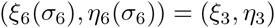. Then, in this case, the relevant equilibria are the ones in the segment corresponding to 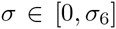. In a similar way, if 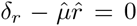, the equilibria 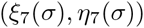 have non negative components for 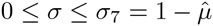. This corresponds to the segment determined by 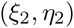 and 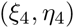.

To study the stability of the equilibrium points we consider the matrix of the linearized system. This matrix has the following form

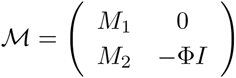

where *I* is the identity of order 4, and

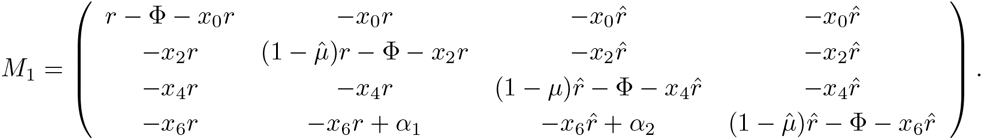

*M*_2_ is not necessary to our purposes. We note that -Φ < 0 is an eigenvalue of *M* with multiplicity 4.

The eigenvalues of *M*_1_ at the non negative equilibria are the following

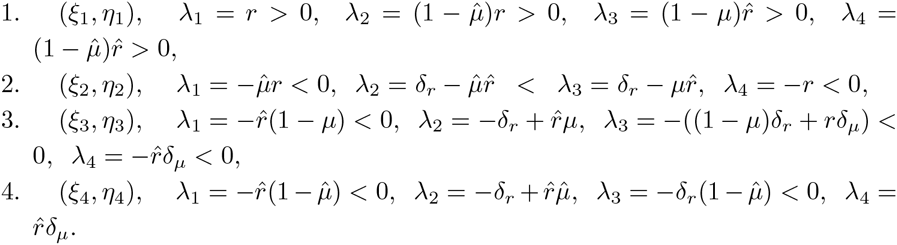

We notice that when 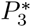 is unstable, 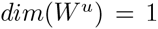. For 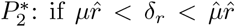, 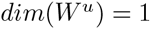; and if 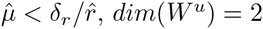.

It is clear that for any given set of admissible parameters, 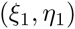 is unstable. Moreover, if 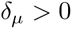, as we are assuming, then 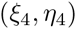 is also unstable.

The stability of 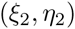 and 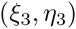 depends on the sign of 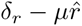.

- If 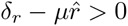 then 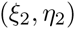 is unstable and 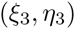 is stable.
- If 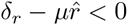 then 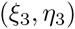 is unstable and 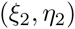 is stable.

That is, from the previous conditions we can derive the bifurcation value involving the transition from tumor cells dominance to tumor clearance. Such a bifurcation value can be written

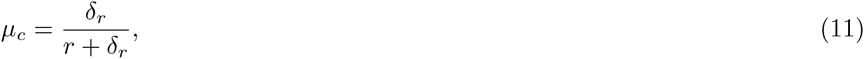

or, alternatively, as

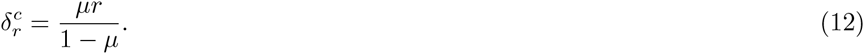

When 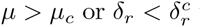 the population of tumor cells will collapse to extinction (see Proposition 1.2. for a detailed explanation of the nature of the bifurcation identified).

We remark that if *δ_µ_* = 0, the equilibrium points 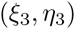 and 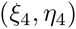 coincide. Moreover if we accept that δµ crosses the value 0, the points interchange stability. However, this is not a biologically meaningful scenario since always *δ_µ_* > 0.

The bifurcation values (11) and (12) involve, when surpassed, an abrupt transition between tumor persistence and tumor clearance. This discontinuous transition is displayed in Fig. 1C. Here we numerically compute the equilibria of the populations at increasing *µ*. In all of our numerical simulations we will consider as initial conditions a tiny population of tumor cells in a healthy tissue with *x*_0_(0) = 0.94. We note that the three neighbors of node 000 do not receive mutational income from 000 because healthy cells are assumed to have mutation rate 0. Hence, we use *x*_1,2,4_(0) = 0.02 as initial conditions. All other variables, which are the lethal phenotypes and sequence 110, are set to *x*_3,5,6,7_(0) = 0. The value of the critical mutation rate in Fig. 1C, is given by *µ_c_* = 1/3, which perfectly matches with our theoretical prediction. Notice that when *µ* < *µ_c_* the equilibria of the lethal populations monotonically increase as mutation rises up, while the equilibria of sequence 100 fastly decreases. The other variable, *x*_6_ also increases, but all populations collapse at the critical mutation rate. Notice that the collapse is discontinuous (see Proposition 1.2. for a detailed explanation of this nature of transition). Panels in Fig. 1D display the time dynamics below and above *µ_c_*.

Figure 2 displays the equilibria in the parameter spaces (*µ*, *δ_µ_*). Here, as we identified in the theoretical calculations, the bifurcation involving tumor extinction does not depend on *δ_µ_*. However, an increase in *δ_µ_* slightly increase the population number of lethal sequence 101. Interestingly, increasing *δ_µ_* involves a decrease in populations 110, while populations with sequence 100 largely increase. This indicates that, below the bifurcation, the increase of genomic instability favors the proliferating tumor cells. Two phase portraits below (Fig. 2B1) and above (Fig. 2B2) the critical mutation rate are also displayed by means of two-dimensional projections in the simplex. In each phase portrait we plot several initial conditions, also including the fixed points living in the simplex for each chosen parameter combination. Below the bifurcation value (with *µ* = 0.27), the fixed points 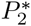 and 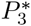 are, respectively, unstable and stable, involving the out competition of the healthy cells by the tumor ones. On the contrary, with *µ* = 0.35 > *µ_c_*, the stability is exchanged between these two distant points, causing the abrupt transition towards tumor extinction. Figure 3 displays the equilibria in the parameter spaces 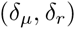. Since the bifurcation value also depends on *δ_r_*, we can see those values of 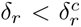 involving tumor clearance and survival of healthy cells. As we previously found, the increase of *δ_µ_* for 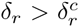 here also favors the population of cells with sequence 100 with higher proliferation rates.

**Figure 2.**
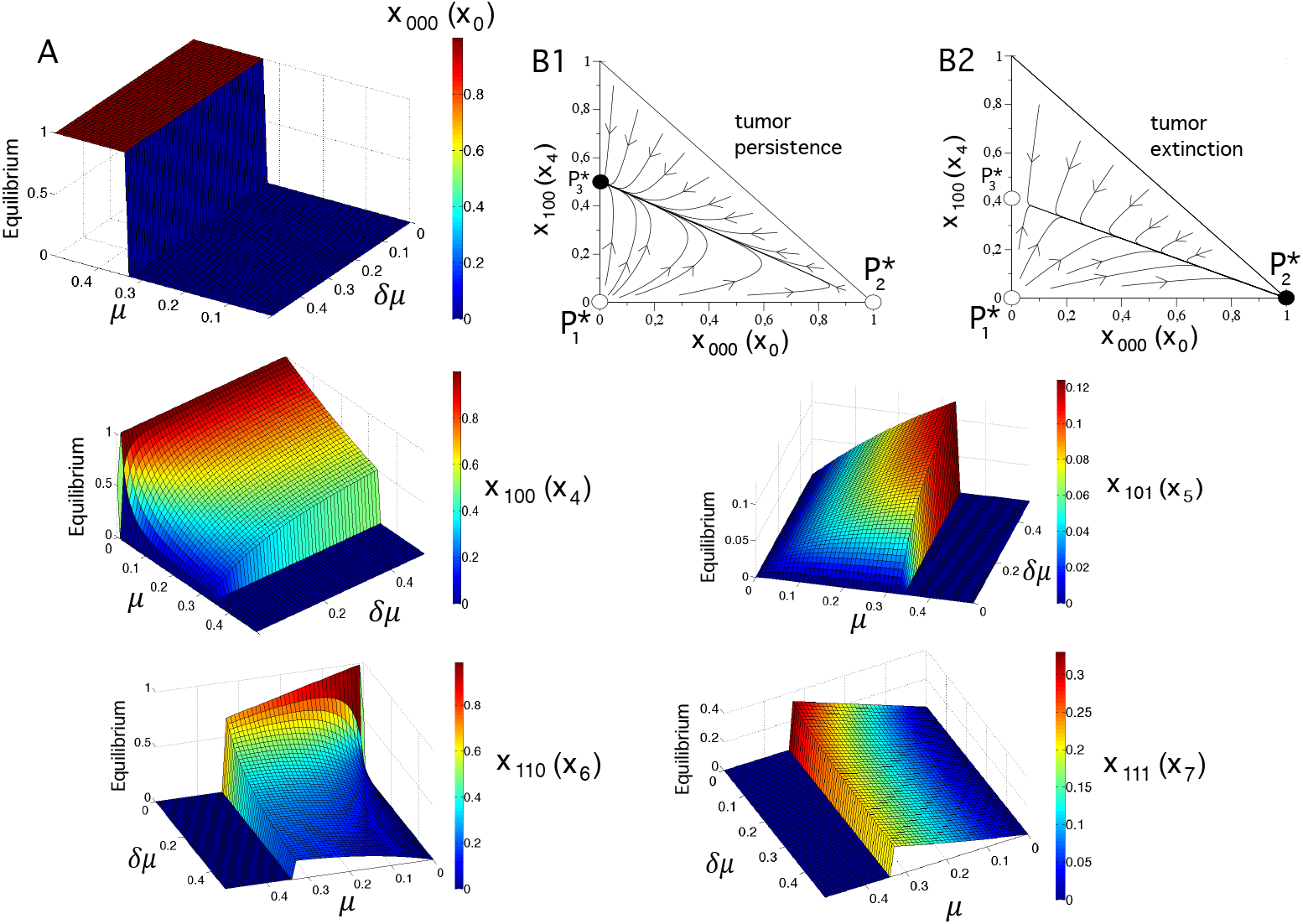
Effect of *µ*, and *δ_µ_* on the equilibrium populations. (A) The surfaces display the equilibrium concentrations of the state variables represented in the parameter space (*δ_µ_*, *µ*), setting *δ_r_* = 0.05. The other variables i.e., *x*_001_(*x*_1_); *x*_010_(*x*_2_); *x*_011_(*x*_3_) have zero population numbers at equilibrium. (B) Dynamics projected on the phase portrait (*x*_0_*,x*_4_), also setting *δ_r_* = 0.05, and *δ_µ_* = 0.3 with Panel 1: *µ* = 0.27; and Panel 2: *µ* = 0.35. Notice that the straight line to which all initial conditions flow corresponds to the heteroclinic connection between fixed points 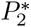 and 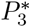. The fixed point 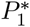 is also shown on the simplex (stable and unstable equilibria are indicated in black and white, respectively). In all the analyses we set *r* = 0.1.

**Figure 3.**
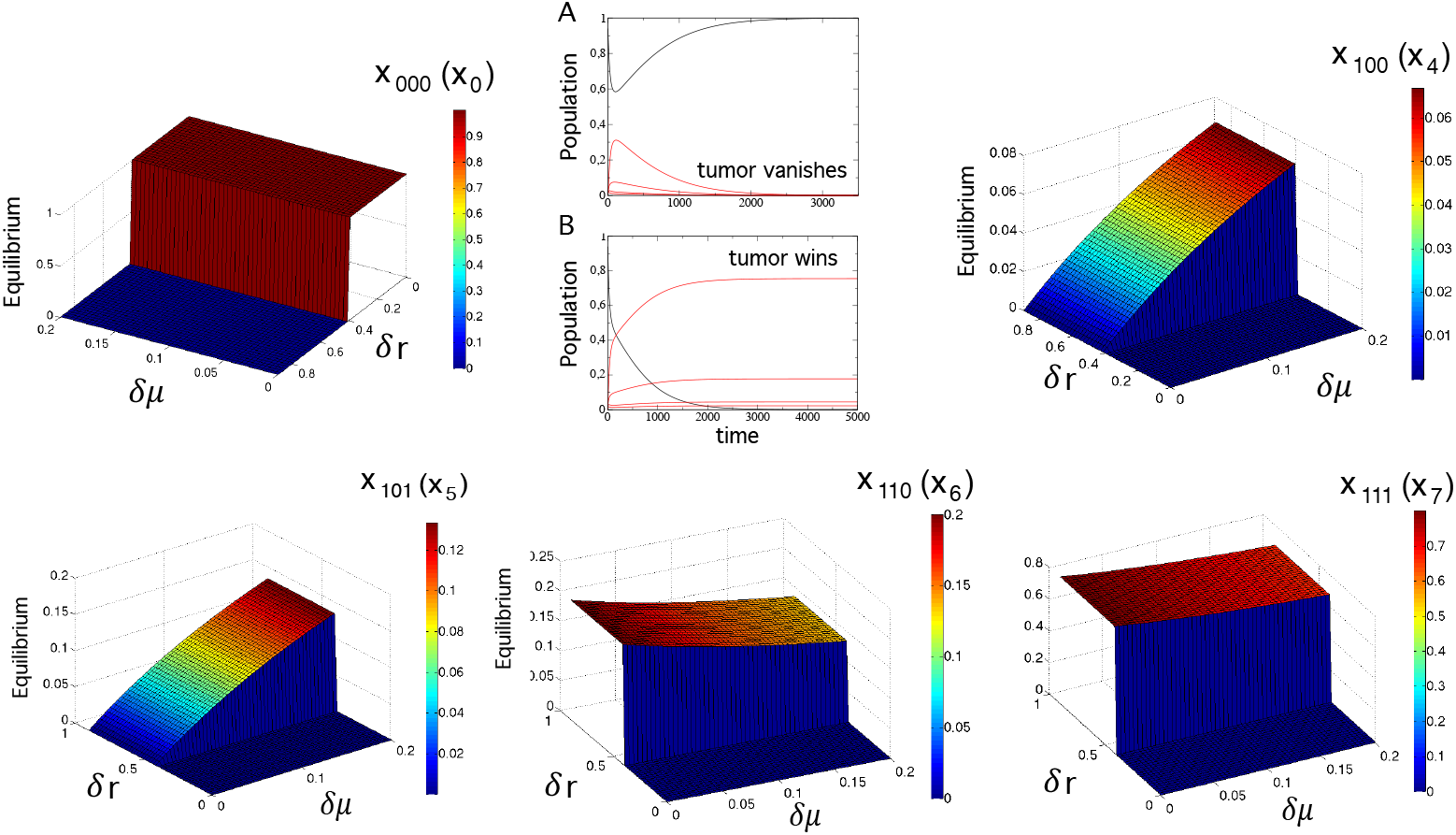
Effect of *δ*_µ_ and *δ_r_* on the equilibrium populations. Here the surfaces also display the equilbrium concentrations of the state variables represented in the parameter space (*δ*µ, *δ_r_*) at high mutation rates i.e., *µ* = 0.8. As in the previous figure, state variables *x*_001_(*x*_1_); *x*_010_(*x*_2_); and *x*_011_(*x*_3_) have zero population numbers. In (A) we display the scenario where healthy cell populations outcompete the tumor population, with *δ_r_* = 0.39, once the critical value of *δ_r_* is crossed. If *δ_r_* is increased, the tumor outcompetes the population of healthy cells, as we show in (B) using *δ_r_* = 0.41. In both panels we used *δ_µ_* = 0.05. In all the analyses we use *r* = 0.1.

### 1.2 Explicit solutions

Now we look for explicit formulas for the solutions of (10). To do so, it is useful to write the system as

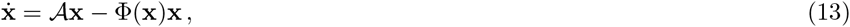

where 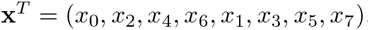,

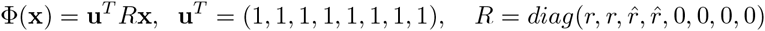

and

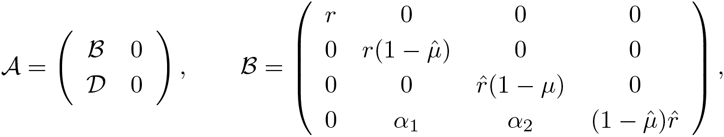

and 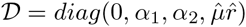.

As usual if we introduce new variables 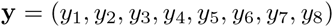 given by

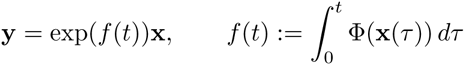

integrating along a solution x(*τ*). Then we obtain a linear constant coefficients system for y

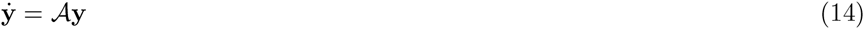

that can be solved easily. The initial conditions are y(0) = x(0). To recover × we use the invariance of 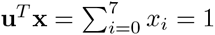 to write

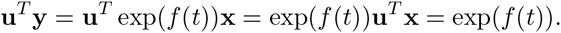

Then,

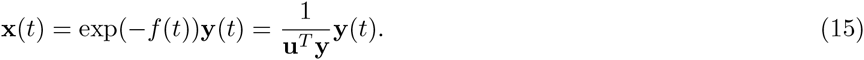

To simplify the notation we introduce

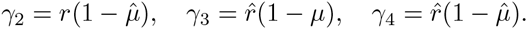

We are assuming that *δ_r_* > 0 and *δ_µ_* > 0. Then γ_2_ ≠ γ_4_ and γ_3_ ≠ γ_4_. The solution of (14), which satisfies 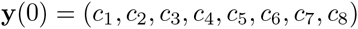, is

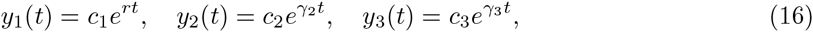

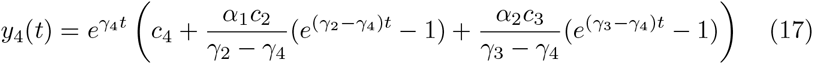

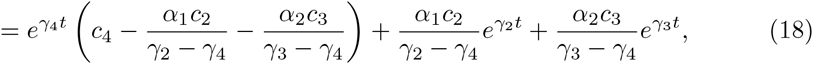

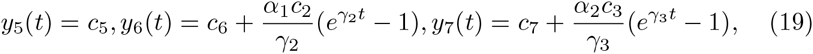

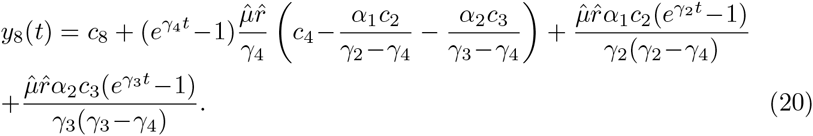

Given some initial conditions 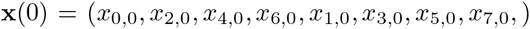, tak-ing into account that 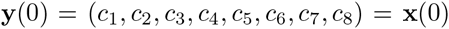 and using (15) we recover the initial variables x, that is

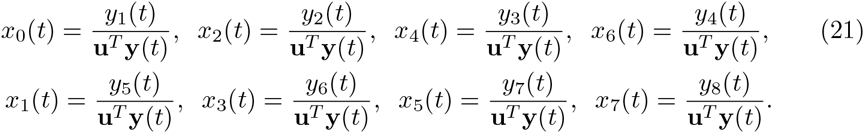

#### Proposition 1.1

*Assume* 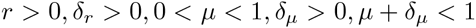 *are fixed values. Let* x(*t*) *be the solution of* (*10*) *such that* 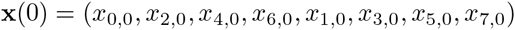, with 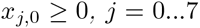, and 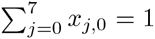. Then for *any* t ≥ 0, all the *components of* x(t) are non *negative. Moreover*

- Assume 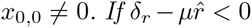 then x(*t*) *tends to the equilibrium point* (*ξ*_2_,*η*_2_) as t goes to *infinity*.
- *Assume* 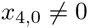. *If* 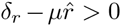 *then* x(*t*) *tends to the equilibrium point* 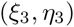 as *t goes to infinity*.
- *Assume* 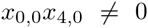. If 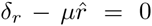 then x(*t*) *tends to the equilibrium* 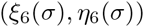*, as t goes to infinity, where*

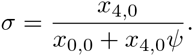

*We recall that ψ has been dened as* 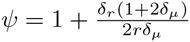.

**Proof** We note that 0 < γ_2_ < γ_4_ < γ_3_. Assume that the initial conditions are non negative. Then it is clear from (16), (17) and (19) that yi(t) ≥ 0 for *i* = 1, 2, 3,4, 5, 6, 7. Moreover using that 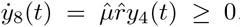, we obtain that *y*_8_(*t*) is an increasing function of *t*. So, if 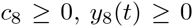. Therefore, 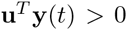 and *x_i_*(*t*) ≥ 0 for all *t* ≥ 0.

- If 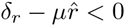, then *r* > γ_3_ and the dominant terms in (16) - (20) are the ones corresponding to *e^rt^*. In particular, if 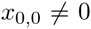, then 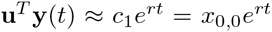 and the solution (21) goes to the equilibrium point 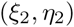 as *t* goes to infinity.
- If 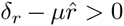, then *r* < γ_3_. If we assume that 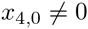 the dominant terms are the ones corresponding to *e*^γ_3_^*t*^^ and

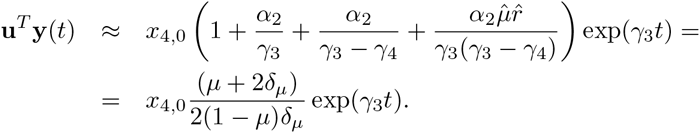

In this case, as *t* goes to infinity, the solutions tend to the equilibrium point 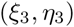.

- If 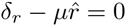, then *r* = γ_3_. Now the dominant terms are

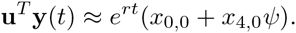

If 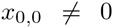 or 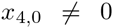, as 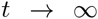 the solution tends to the point 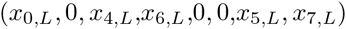 where

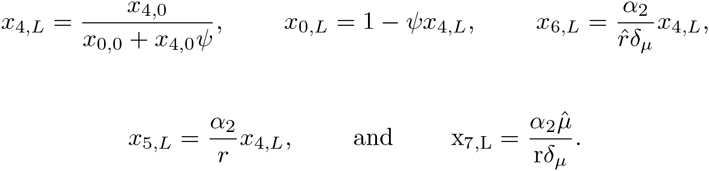

That is, the limit point is 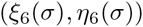, with 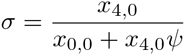.

#### Proposition 1.2

*Assume that at t = 0 we start at a point of the segment joining the points 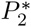 and 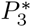 of the form 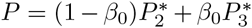. Then the solution remains on that segment for all time, and at time t it is located in 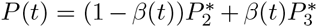, where*

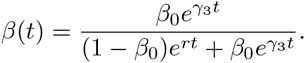

The proof follows immediately from the expressions of the solutions (16-20). We note that this result is valid for all values of the parameters.

Hence, for values of *µ* close to *µ_c_* the situation is as follows: For *µ* > *µ_c_* the solution 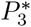 is unstable, with an unstable 1-dimensional manifold and one of the branches tends to the stable point 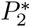. This is a heteroclinic connection. For *µ* < *µ_c_* the situation is reversed: the heteroclinic connection goes from 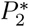 (unstable) to 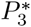 (stable). For the critical value *µ_c_* all the points on the line passing through 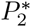 and 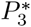 are fixed. Furthermore, for *µ* close to *µ_c_*, the full line is attracting. We can talk about a *trans-heteroclinic* bifurcation.

As for *µ* close to *µ_c_* the eigenvalues associated to the line joining 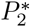 and 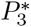 are close to zero, and all the other eigenvalues at these points are negative, arbitrary initial conditions (except a set of zero measure) tend to that line. So, we can reduce the study to the dynamics to a single line. A simple model of the *trans-heteroclinic* bifurcation is as follows.

Assume that a differential equation in dimension one depends on a parameter *α* and for any value of the parameter the points *x*_0_ = 0, *x*_1_ = 1 are fixed. Consider the model equation

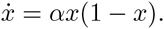

Then, for *α* > 0 points between *x*_0_ and *x*_1_ move towards *x*_1_, while for *α* < 0 they move towards x_0_. For *α* = 0 all the points on the line are fixed. This is exactly a model of what happens in the present system.

### 1.3 Bifurcations and transients: Implications and potential therapies

The previous calculations and numerical simulations revealed a catastrophic transition separating tumor persistence from extinction. The bifurcation value has been shown to depend on both mutation and proliferation rates of tumor cells. Here we will focus on these two parameters seeking for the effects of their manipulation with potential therapies. Several strategies to directly target cancer cells (targeted cancer therapies) have been discussed. For instance, DNA nano clews containing drugs could be used for drug delivery inside tumor cells [42] (see below and Conclusions Section). In Fig. 4A we display a bifurcation diagram increasing mutation rates of tumor cells for 5 different values of the proliferation rate of tumor cells, *δ_r_*. Notice that the decrease of the replicating tumor populations (*x*_4_ + *x*_6_, red line) is linear at increasing mutation. Interestingly, when the increase in the proliferation rate is tiny (e.g., *δ_r_* = 0.01), the critical mutation value, *µ_c_*, becomes very small. This result suggests that possible therapies combining both mutagenic and cytotoxic drugs may allow to achieve the abrupt bifurcation more easily. For larger values of *δ_r_* the critical mutation rate largely increases. This would involve higher concentrations of mutagenic drugs or longer exposures to such drugs (see Conclusions Section).

**Figure 4.**
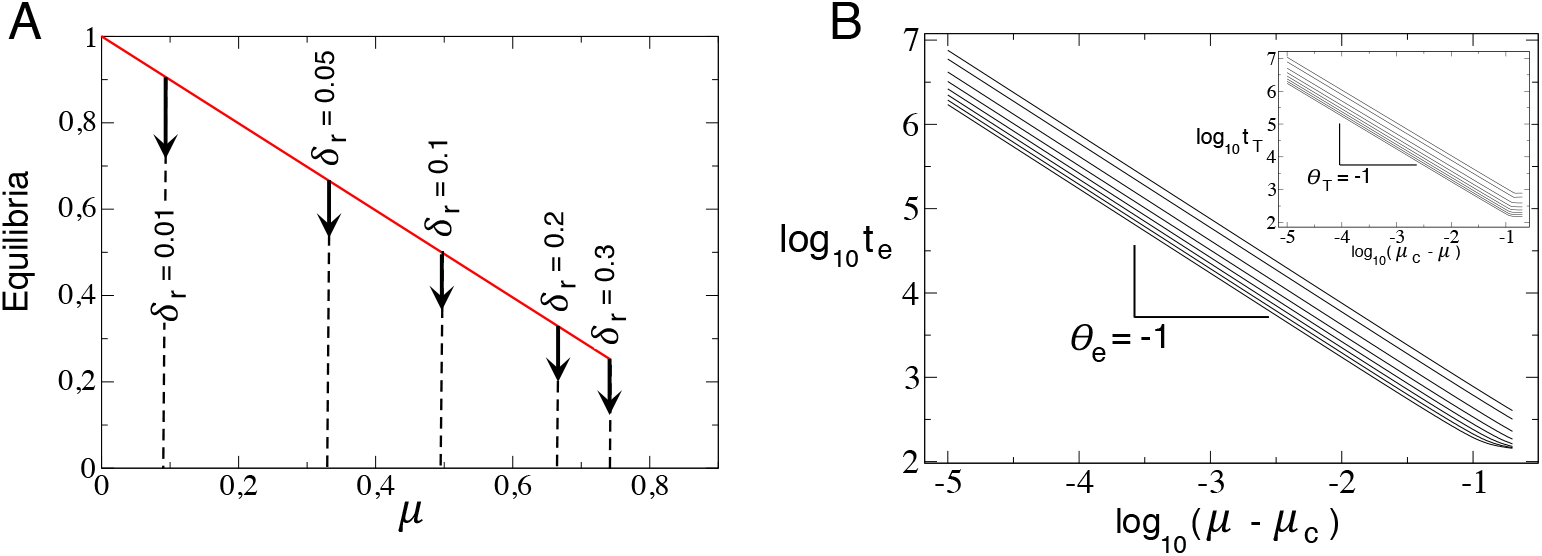
Catastrophic shifts of tumor cell populations and transient times near bifurcation threshold. (A) Abrupt extinction of tumor cell populations (represented as 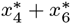, red i.e., not considering lethal phenotypes). The vertical dashed black lines represent the critical mutation rates *µ_c_* given by Eq. (11) for (from right to left) decreasing values of *δ_r_*. (B) Times to tumor extinction, t_e_, as mutation rate *µ* increases above the critical mutation value *µ_c_* for (from top to down): *δ_r_* = 0.01; *δ_r_* = 0.05; *δ_r_* = 0.1; *δ_r_* = 0.2; *δ_r_* = 0.3; *δ_r_* = 0.4; *δ_r_* = 0.5; *δ_r_* = 0.6; *δ_r_* = 0.7. Notice that t_e_ displays a power-law dependence after bifurcation threshold: t_e_ ~ (µ — *µ_c_*)^θe^ with *θ_e_* = —1. The inset displays the time it takes to the system to achieve the fixed point Pg (where the tumor wins) below bifurcation threshold (here we used the same increasing values of δ*_r_* (also from top to down) than in the big plot). Notice that the same exponent is found below bifurcation threshold, i.e., t_T_ ~ (µ_c_ — *µ*)*^θt^, θ_t_* = —1. In both panels we use *r* = 0.1 and *δ_µ_* = 0.1 (we notice, however, that the results of this figure do not depend on *δ_µ_*).

Another interesting point that we can address with the model are the transients towards both fixed points 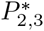 as we approach to the bifurcation value (from below and above), and their dependence on *δ_r_*. Here, recall that with *µ* < *µ_c_* the fixed point 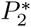 is unstable and 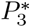 is stable. Contrarily, with *µ* > *µ_c_* the fixed points 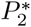 and 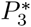 are, respectively, stable and unstable. The transients are computed as the time a given trajectory *ϕ_t_* spends to achieve a distance 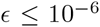 to the fixed point 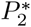 or 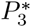, depending if or *µ* > *µ_c_*. Here we will also use as initial conditions 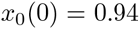, 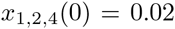, and 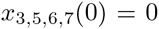. The results are displayed in Fig. 4B. Here we numerically computed the transient times near but above *µ_c_* (big panel) and near but below *µ_c_* (inset). The numerical results reveal a power-law dependence of the transient times with the distance to the bifurcation value. For example, above bifurcation threshold, the time that the tumor cells spend towards extinction, *t_e_* (trajectories are attracted by 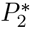), scales according to 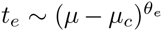, with *θ_e_* = —1. This scaling law (see below for the mathematical derivation of this exponent) indicates that with a mutation rate extremely close to *µ_c_* the system spends a huge amount of time before tumor extinction. However, a tiny value increase beyond this critical value the system drastically reduces the time-to-extinction, meaning that responses time would be extremely fast after drug treatment. In this scenario, it is worth to mention that tumor populations with higher proliferation rates would become extinct more rapidly, although the critical mutation rate is also much higher (notice that Fig. 1B gives information about the transients as mutation is changed from the bifurcation value, however, the critical mutation value grows as *δ_r_* grows, see Fig. 1A). For instance, the time-to-extinction with 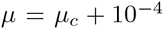 is 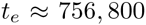 for 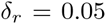; and 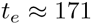 for 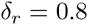. If we increase mutation this time is drastically reduced: the time-to-extinction with 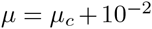 is 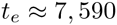 for 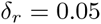; and 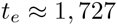 for *δ_r_* = 0.8 (times are in arbitrary units).

The inset in Fig. 1B displays the same information now tuning mutation up to *µ_c_* but from below. This means that the orbits are attracted by the fixed point 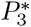, which involves tumor persistence and extinction of the healthy cell populations. Here, the time the tumor spends to outcompete the population of healthy, *t_T_* cells scales according to 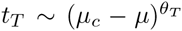, with 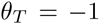. Here, the higher the value of *δ_r_*, the shorter the time the tumor spends to achieve equilibrium 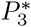. This means that increasing mutation rate close, but below *µ_c_* would involve a fast tumor growth impairing the healthy population. This strategy would not be so dangerous for slowly-replicating tumors, which also have a low value of *µ_c_*.

In the following lines we will derive the scaling exponent previously identified numerically. As it was mentioned in the previous section, for *µ* close to *µ_c_* the dynamics tends first to approach the segment between 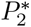 and 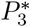. For concreteness assume that it approaches a point P of the form 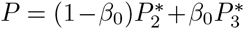. Then the subsequent evolution is of the form 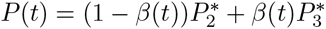, where

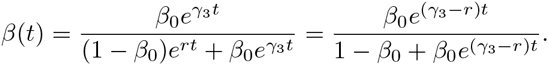

Note that 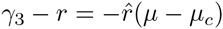.

Hence, for 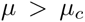 as 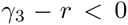 the second term in the denominator becomes negligible in front of the first one after some time. It remains, essentially, 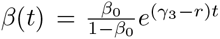 hich tends to 0 as time increases. That is, the point tends to 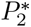. If we want to stop at a distance *d* from 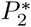 we put 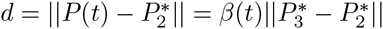 and denoting as *C* the value of 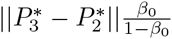 we obtain 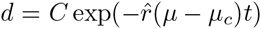, where we have used the expression of 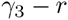 given above.

Taking logarithms twice in the last equality one has 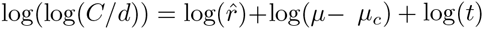.

For fixed values of *r,δ_r_,d* and a given initial value *β*_0_ this is the equation of a straight line in the 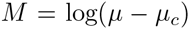 and 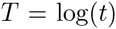 variables, with slope —1; *M + T = S*, where *S* denotes 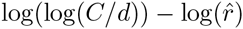.

Hence, *t* is of the order of magnitude of 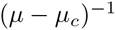. We also check that increasing *δ_r_* and, hence, increasing 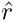, the value of *S* decreases and the line with slope -1 is going a little bit down, as seen in Fig. 4B.

The same reasoning applies if 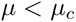 looking for the approach to 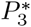.

Anyway, we should note that for values of 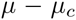 relatively large one can have relevant variations in the value of the parameter *β*_0_ of the point of the segment between 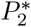 and 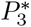 that we approach at the beginning of the evolution. This is clearly reflected on the figure when the values of 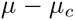 exceeds 0.1. Furthermore, this effect is more visible in the case 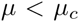, when approaching 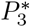. This is due to the fact that the initial point, with first component 0.94, is much closer to 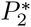. The effect of this first component is also reflected on the fact that, for a fixed value of *δ_r_*, the time to be at distance *d* from 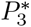 with 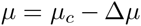 is larger than the time to be at distance *d* from 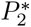 with 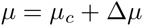.

## Conclusions

Most of the transitions and bifurcations identified in deterministic quasispecies models towards the so-called error-threshold are given by smooth, continuous shifts [38-41]. Moreover, previous mathematical models [15,16,26,27,33] applied to cancer dynamics have also revealed smooth transitions towards the impairment of cancer populations. Here, we identified a novel bifurcation named as *trans-heteroclinic bifurcation*, which involves the stability exchange between two equilibria placed far away from each other in phase space. These conditions lead to a non-smooth, i.e., catastrophic, transition governing the extinction of tumor cell populations. As a difference from transcritical or saddle-node bifurcations of fixed points or periodic orbits, the two equilibria in our system do not collide in phase space. We notice that the bifurcation found in our model is not very standard and involves a transcritical-like bifurcation with a discontinuous shift as a given control parameter crosses its bifurcation value. As a difference from a transcritical bifurcation, the fixed points do not collide when interchanging stability, thus giving place to an abrupt transition

The bifurcation value in our system depends on the mutation rate of tumor cells and on their proliferation increase. This finding can give clues about how a proper therapy may affect the tumor behaviour. In this sense, the so-called targeted cancer therapies are being designed to deliver drugs inside cancer cells in a specific manner. For instance, DNA nano-capsules storing anticancer drugs [42]. Roughly, such nano capsules could attach to the receptors of the tumor cells, being internalised inside them and delivering the drugs after their self-degradation inside the cancer cell lysosomes. For this particular case, recent essays have studied the release dynamics of the anticancer drug DOX (doxorubicin) in an acidic medium *in vitro* [42]. DOX is known to act on macromolecular synthesis, inhibiting the progression of the enzyme topoisomerase II during DNA replication. Hence, DOX has a cytotoxic effect stoping the process of replication reducing cancer cells proliferation. Another side effect of DOX is the increase of free radical production, which could involve increased mutagenesis in such cells. In this sense, it is known that oxidative damage can increase the mutation frequency by an average 4.3-fold [43].

The results reported here highlight the importance of combination therapies using both mutagenic and cytotoxic drugs, such as DOX. Similarly, other mutagenic drugs such as etoposide, mitoxantrone and teniposide (other topoisomerase inhibitors) could be applied to cancer cells by means of the DNA nano capsules, in combination with cytotoxic drugs such as maytansine [44], vinorelbine, vindesine, and vinflunine [45] (which block cell division by inhibiting the assembly of microtubules). The combination of these two drugs could, according to our results, drive the system towards parameter values causing the abrupt collapse of the tumor populations.

## Competing interests

The authors declare that they have no competing interests.

## Author’s contributions

J.S and R.V.S. built the mathematical model. C.S. and R.M. carried out the analysis of the mathematical model and the related theoretical calculations. J.S. and C.S. produced the numerical data. J.S., C.S., R.M. analyzed the numerical data. All authors wrote the manuscript.

## Acknowledgements

We thank the members of the Complex Systems Lab for their helpful comments, as well as the members of the Instute of Evolutionary Biology (IBE) for their critics and comments. The authors acknowledge the computing facilities of the Dynamical Systems Group (Universitat de Barcelona). This work was partially funded by the Botín Foundation (JS, RVS), by the Spanish grant FIS2012-39288 (RVS), by the Santa Fe Institute (RVS), by the Spanish Secretaria de Estado de Investigación, Desarrollo e Innovación grants MTM2013-41168-P (CS, RM), and by grant 2014-SGR-1145 from the Catalan government (CS, RM).

